# A new device for continuous non-invasive measurements of leaf water content using NIR-transmission allowing dynamic tracking of water budgets

**DOI:** 10.1101/2022.05.06.490892

**Authors:** Hartmut Kaiser

## Abstract

Leaf water content (LWC) permanently fluctuates under variable transpiration rate and sap flow and influences e.g. stomatal responses and osmotic adjustment of plant cells. Continuous recordings of LWC are therefore central for the investigation of the regulatory networks stabilizing leaf hydration. Available measurement methods, however, either influence local hydration, interfere with the local leaf micro-environment or cannot easily be combined with other techniques. To overcome these limitations a non-invasive sensor was developed which uses light transmission in the NIR range for precise continuous recordings of LWC. For LWC measurements the transmission ratio of two NIR wavelengths was recorded using a leaf-specific calibration. Pulsed measurement beams enabled measurements under ambient light conditions. The contact-free sensor allows miniaturization and can be integrated into many different experimental settings. Example measurements of LWC during disturbances and recoveries of leaf water balance show the high precision and temporal resolution of the LWC sensor and demonstrate possible method combinations. Simultaneous measurements of LWC and transpiration allows to calculate petiole influx informing about the dynamic leaf water balance. With simultaneous measurements of stomatal apertures the relevant stomatal and hydraulic processes are covered, allowing insights into dynamic properties of the involved positive and negative feed-back loops.

## Introduction

A sufficient and stable hydration of cells is essential for aerial plant organs to survive in the often desiccating atmosphere. In the leaves of vascular plants various mechanisms have evolved to provide sufficient water uptake while restricting and regulating transpirational water loss to ensure a stable leaf water content which not only stays within safe margins for survival, but is optimal for cellular processes like photosynthesis. Maintaining a stable hydration under any combination of permanently fluctuating often adverse environmental influences requires the interaction of various sensing and regulating mechanisms tightly controlling both the uptake of water to leaves and transpirational losses through stomatal pores. Tissue water content can thus be seen as one of the most tightly controlled physical properties of vascular plants. Many of the mechanisms regulating leaf water content form negative feedback loops whereby deviations of LWC or related physicochemical properties are sensed and transformed into short, mid or long term responses aimed to recover the target state of water content. Responses range from short term adjustments (e.g. stomatal responses) over mid-term osmotic adjustments to long term growth responses leading to morphological adaptations (Osakabe et al. 2014; Bielach et al. 2017; Buckley 2019; Chérel and Gaillard 2019). Water status in these feedback loops is both the controlled condition and serves as the input to the homoiostatic feedback-loops. The most important and effective way to regulate leaf water content is the control of leaf transpiration by osmotically driven changes of guard cell turgor and hence pore-width. The mechanism by which guard cells are triggered to respond to changing LWC however is not yet fully understood despite of a huge amount of studies published on this topic over the last decades.

Much of the research revolving around stomatal responses and their role in controlling leaf hydration use measurements of stomatal responses and transpiration by leaf gas exchange at a sufficient time resolution and accuracy. Unfortunately, the property to be controlled, namely LWC, cannot easily be measured simultaneously with equally high accuracy and temporal resolution which impedes the analysis of the dynamics of feedback control. The classical approach to measure water potential in a pressure bomb (Scholander et al. 1965) is destructive and non-continuous. However, several nondestructive methods to sample LWC or other related measures (turgor, water potential, sap flow) have been developed (Zimmermann et al. 2008; Martinez et al. 2011; Davis et al. 2012; Defraeye et al. 2014; Dadshani et al. 2015; Baldacci et al. 2017; Fariñas et al. 2019). Each of these methods has its experimental benefits, but also comes with specific drawbacks. Some measurements have a poor time-resolution (sap flow recording), affect water status by impeding transpiration and/or changing the energy balance (e.g. psychrometric methods and those methods requiring continuous clamping of leaves) or cannot easily be used within cuvettes with full control of the leaf-micro-environment. For contactless measurements of LWC reflection or absorption of acoustic or electromagnetic waves can be used. Noninvasive ultrasonic resonance spectroscopy (Sancho-Knapik et al. 2016; Fariñas et al. 2019) has proven to yield signals closely related to leaf hydration in many species. While this approach is based (indirectly) on the covariation of leaf mechanic properties with water content, measuring the interaction of electromagnetic waves of different frequences (Sancho-Knapik et al. 2011; Browne et al. 2020) with water molecules promises to offer a more direct measure of water content. Leaf transmission and reflection spectra in the visible light and infrared range (Jacquemoud & Baret, 1990) have widely been employed for contactless determination of leaf and vegetation water content (Thomas *et al*., 1971; Inoue *et al*., 1993; Cozzolino, 2017; Braga *et al*., 2021). This approach has gained importance especially in satellite and aerial vehicle based remote sensing. While absorption by liquid water is an important determinant of IR absorbance and reflection spectra of leaves, a straightforward derivation of LWC from spectral measurements is hindered by interfering effects of e.g. dry matter content, metabolite composition and leaf structural parameters which can widely vary between and even within species. Various solutions to this problem have been proposed and successfully applied: Inclusion of more spectral bands up to continuous spectra, development of various indices (Jiang et al. 2018; Braga et al. 2021; Li et al. 2021), inversion of leaf optical models (Jacquemoud and Baret 1990; Féret et al. 2019) and neural networks (Conejo et al. 2015; Koirala et al. 2020; Braga et al. 2021) successfully aid in deriving specific leaf properties in the presence of confounding variation in overlapping spectral effects.

The approach presented here, however, attempts to simplify the estimation of LWC from spectral measurements by performing calibrations for individual leaves. In this way, even limited spectral information from key spectral ranges strongly influenced by liquid water could be sufficient to get precise measurements of LWC. The separation from non-water-related confounding leaf optical effects is assigned to the calibration procedure which is performed on and valid only for the measured leaf. By measuring in only two spectral IR ranges (Seelig et al. 2009) a simple and miniaturized measurement setup based on two LED’s and one photodiode is possible. Using this approach a non-invasive, precise and continuous optical method to record of LWC is presented which puts little constraint on simultaneously occurring other measurements or the choice and control of experimental conditions. Example measurements demonstrate how the combination of this measurement of LWC with other methods can yield better insight into leaf water budgets and the regulatory processes behind leaf water homeostasis.

## Materials and Method

### Principle of the water content sensor

The transmission of light through leaf tissues depends on specular reflection, internal reflection, scattering and absorption (Govaerts et al. 1996). Varying water content causes changes in the absorption spectrum especially in the distinctive absorption bands of liquid water. In the presented device, the infrared transmission at 1450 nm, a local maximum of the water absorption spectrum, is used as an indicator of LWC. Initial experiments with a first prototype using only the 1450 nm spectral band confirmed a strong response to LWC. However, the leaf clamped by an annular clamp around the measured area, moved slightly due to turgor changes which strongly influenced the IR transmission signal. These movements were evident as a changing focus position of the microscope when simultaneously observing the lower epidermis. Leaf movements change the distance between the light sensing photodiode and the leaf tissue diffusing the measurement beam and therefore strongly influence incident irradiation on the sensor surface. A firmer fixation of the leaf at the spot of measurement, however, is undesirable, as it would inhibit transpiration and thus disturb leaf water relations. This problem was overcome by using a ratiometric measurement principle with an additional beam at 1050 nm with little absorption by liquid water. The 1450 nm and 1050 nm bands were chosen, because they represent two bands which are differentially affected by leaf water content and because both spectral ranges fall into the sensitivity range of InGaAs photodiodes which allowed for a simple setup for transmission measurements basically consisting of only one photodiode and two LED’s. Errors due to leaf movements should affect both measurement wavelengths similarly and be canceled out in the transmission ratio. The same reasoning applies for other errors in the optical and electronical signal chain, which affect both beams independently of their wavelength.

The ratiometric measurement was accomplished by measuring alternately the light transmission of the light emitted by 1450 nm and 1050 nm LED’s (LED1450-03, Roithner Laser, Wien, Austria; LED1050E Thorlabs, Munich, Germany) with an InGAaS photodiode (LAPD-1-09-17-TO46, Roithner Laser, Vienna, Austria). The measuring light was pulsed at 1,2 kHz and photodiode output amplified phase-sensitive with a lock-in-amplifier to allow measurements irrespective of ambient radiation (Fig 1). The optical assembly consists of the two LED’s (pointing to the upper leaf surface at an angle of 45° thus avoiding shading of the measured area when using perpendicular illumination. Radiant power density as estimated from manufacturer data for the LED’s when driven with a current of 20mA in a 50% duty cycle and an illuminated area of 0,5 cm^2^ is lower than 14 W/m^2^, which was assumed to be negligible (approx. 1.4% of full sunlight radiant density). The beams of the LED’s were homogenized by a hexagonal homogenizing light pipe (#63-088, Edmund Optics, Barrington, U.S.) to ensure identical and homogenous illumination for both wavelengths. The photodiode placed 5-10mm below the illuminated area (Fig. 1) picked up the transmitted diffuse light of the LED’s.

**Fig. 1.**
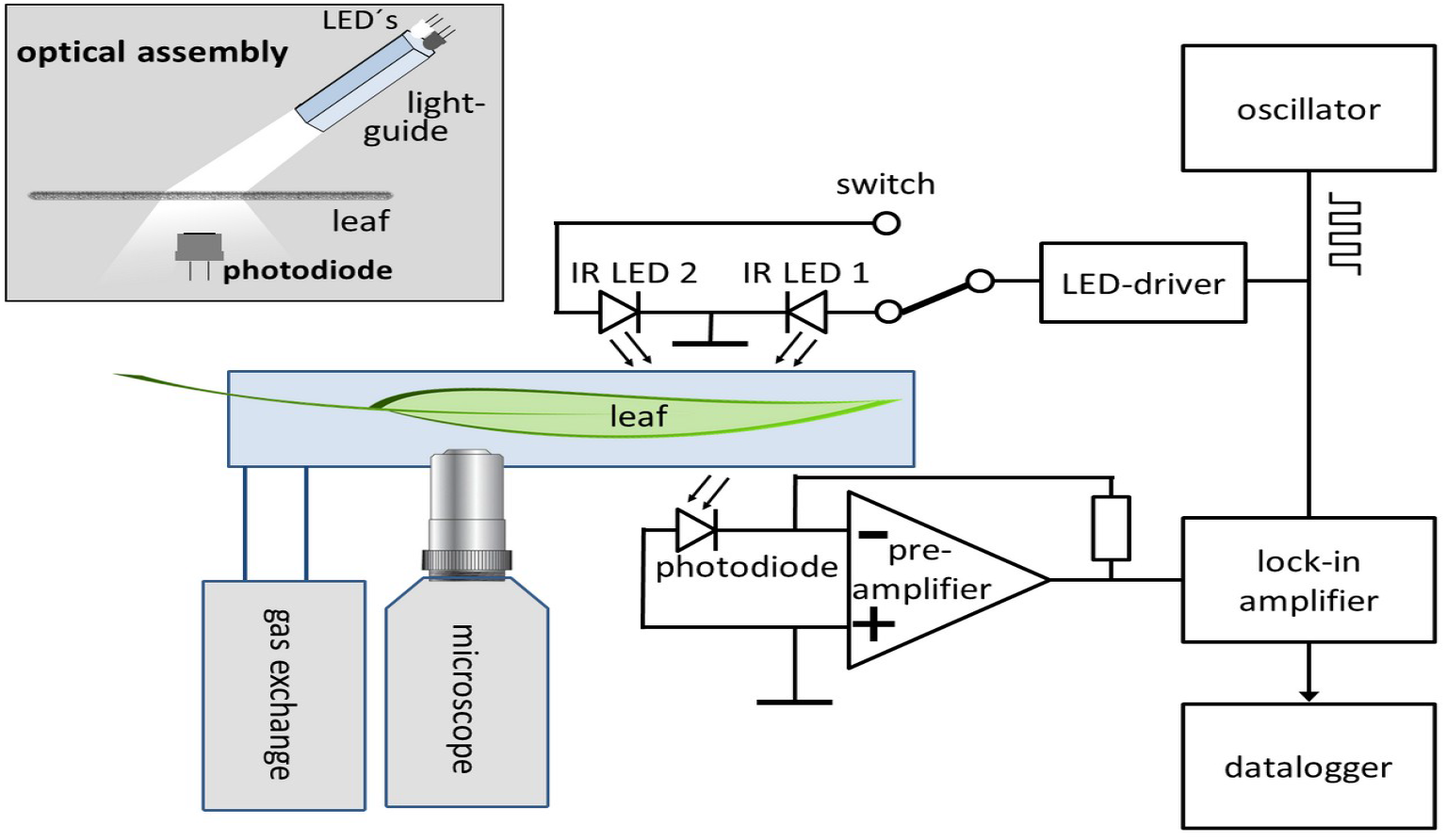
Measuring principle, simplified schematic circuit and optical assembly of the device for the measurement of leaf water content integrated into a cuvette system with simultaneous microscopical observation of stomata.

In order to standardize the I1450/I1050 ratio signal across measurements, the raw signals were divided by the signal obtained using a wavelength independent diffusive standard made of a matte glass slide with scattering properties similar to a typical leaf. This adjustment procedure is performed every time the measurement setup has changed (e.g. changed distance, angle, LED-currents) and warrants comparable I1450/I1050 ratios between measurements.

### Calibration

Accurate calculation of LWC from adjusted I1450/I1050 requires a calibration which is dependent on among others leaf structural parameters and therefore specific for each observed leaf or at least specific for the observed species and growing condition. Calibration can be performed in three different ways. Up to now calibrations mostly were performed after the experiment specific for each measured leaf by simultaneously recording LWC with an independent method and I1450/I1050 ratio. This can be done either by detaching the leaf and weighing the entire leaf clamp together with the leaf on a scale while continuing to measure IR-transmission, which of course requires precautions to keep the force exerted by the cable constant. Another way to calibrate is only possible if the sensor is installed in a gas exchange chamber for continuous transpiration measurements: After cutting the petiole with the leaf still enclosed in the cuvette I1450/1050 ratio and transpiration are measured simultaneously for about 0.5-1 hours. The accumulated transpirational water loss for each measured infrared transmission ratio during this period is finally added to the final LWC determined by weighing fresh leaf weight at the end of the leaf drying period in the cuvette and dry weight after complete drying in an oven. As most of the experiments up to now have been performed in a cuvette, this procedure was the usual way to obtain an accurate per leaf calibration.

Another method of calibration is applicable when per leaf calibration by the balance method and the transpiration based method are not possible. Regression on accumulated data of several leaves of a species or variety under defined growing concitions can result in a calibration specific for a batch of plants in an experiment (fig. 2). To demonstrate such calibrations and their associated measurement uncertainties, leaves of *Vicia faba* plants from two growing conditions (outdoors in pots or hydroponically in a climate chamber) and of *Zea mays* grown in pots in a greenhouse were detached in the fully turgid state, and either immediately weighed and IR transmission ratio measured with the LWC sensor or left to dry under room conditions for c. 10 min before measurement. The obtained relationships between the IR transmission ratio followed a linear relationships as long as LWC was above the wilting point. As the sensor was only used within this range a linear calibration was sufficient and could be used to calculate leaf water contents from IR transmission ratios. The calibration lines differed in slope and offset between species and growing conditions, showing that not only each species requires an individual calibration, but also that growing conditions can substantially change the relation between the IR transmisson ratio and LWC. After calibration for a certain variety under the experimental growing conditions, a measurement uncertainty (experimental standard deviation) of between 6 to 9 g m^-2^ was achieved, which is between 3 to 6% of the total leaf water content.

**Fig. 2.**
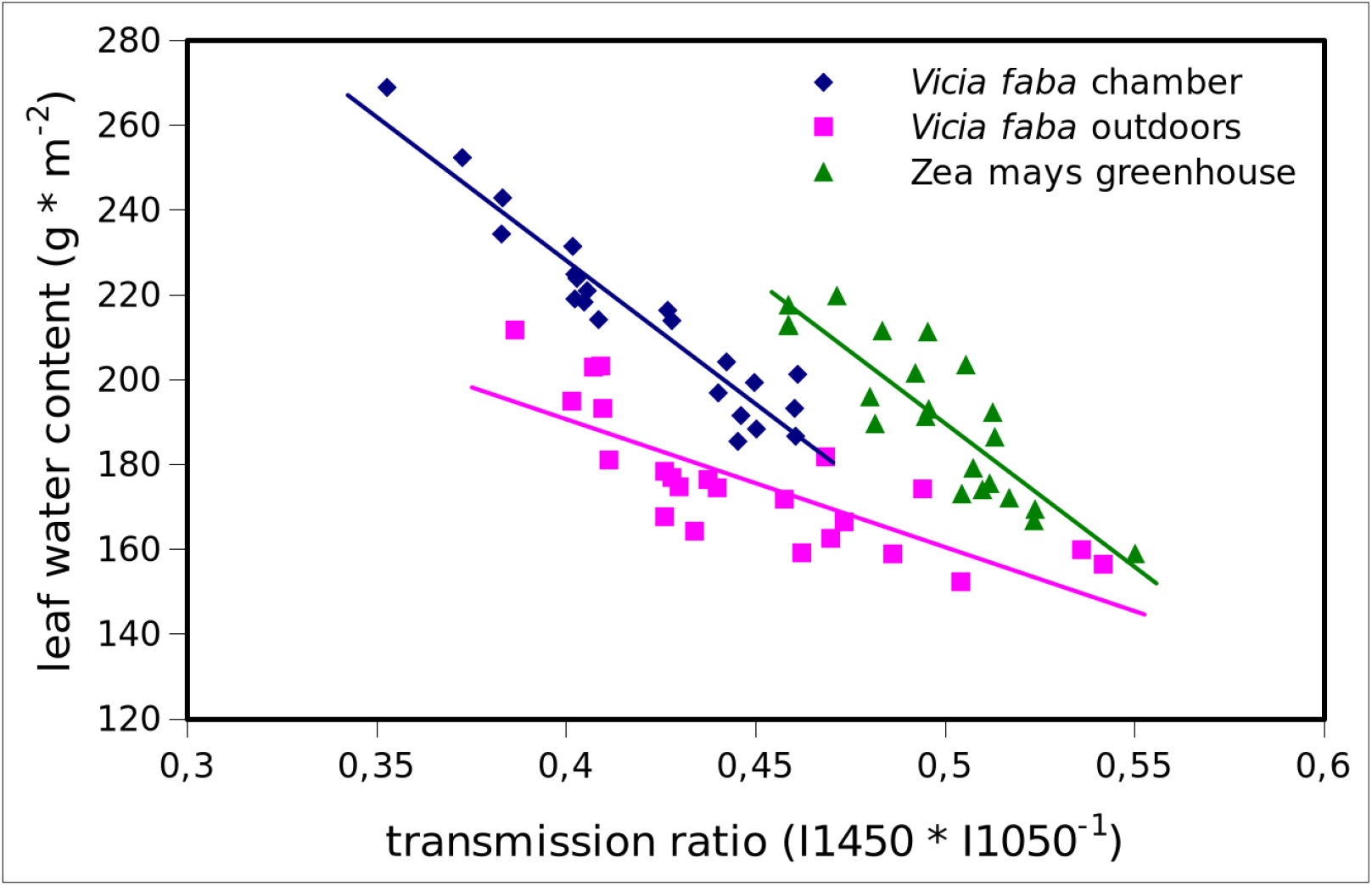
Calibrations of the leaf water content sensor for leaves of Vicia faba grown hydroponically in climate chamber, Vicia faba grown outdoors in Mitscherlich-pots and Zea mays plants grown in a greenhouse. On 22 detached leaves of each species resp. treatment leaf water content was measured gravimetrically simultaneously with the transmission ratio I1450/I1050. To induce variation in LWC, 11 of the leaves were measured immediately after cutting in the fully turgescent state and 11 after c. 10 minutes of passive drying under c. 22°C and 50% rH.

### Quality of the measurement

One possible drawback of the ratiometric approach is the multiplication of noise from the two channels. The final signal-to-noise ratio, however, remains fully sufficient (Fig 3) with an error margin of ±0.0155 g*m^-2^ when using oversampling and averaging to a 20s interval. To put this into perspective this signal noise amounts to about 0,04% of the amplitude of changes in LWC observed during a drought experiment leading to non lethal wilting (fig. 7). This high precision together with the continuous nature of this measurement allowed to discriminate changes in LWC after a change in conditions already after 20 to 30 seconds (figs 4-6).

**Fig. 3.**
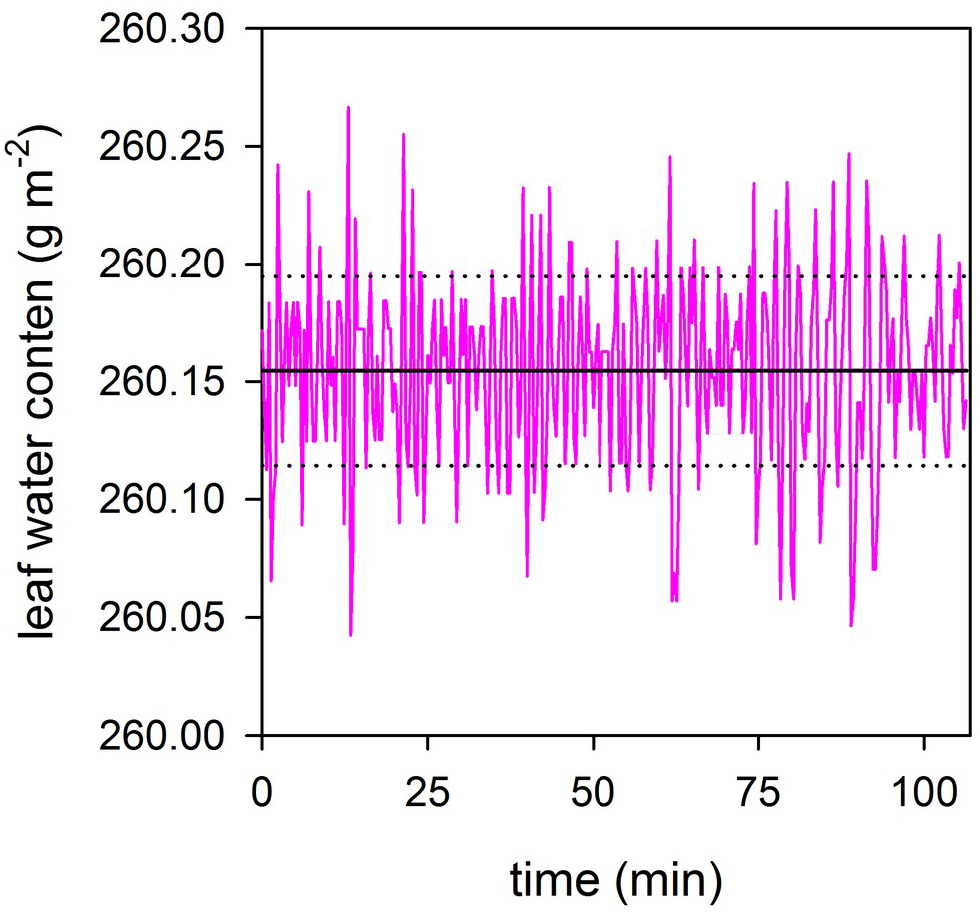
Demonstration of signal to noise ratio (SNR). The measurement was performed on an attached leaf of Vicia faba during a period of stable leaf water content with a running average applied over 30 sec. Dotted lines indicate ±standard deviation.

**Fig. 4.**
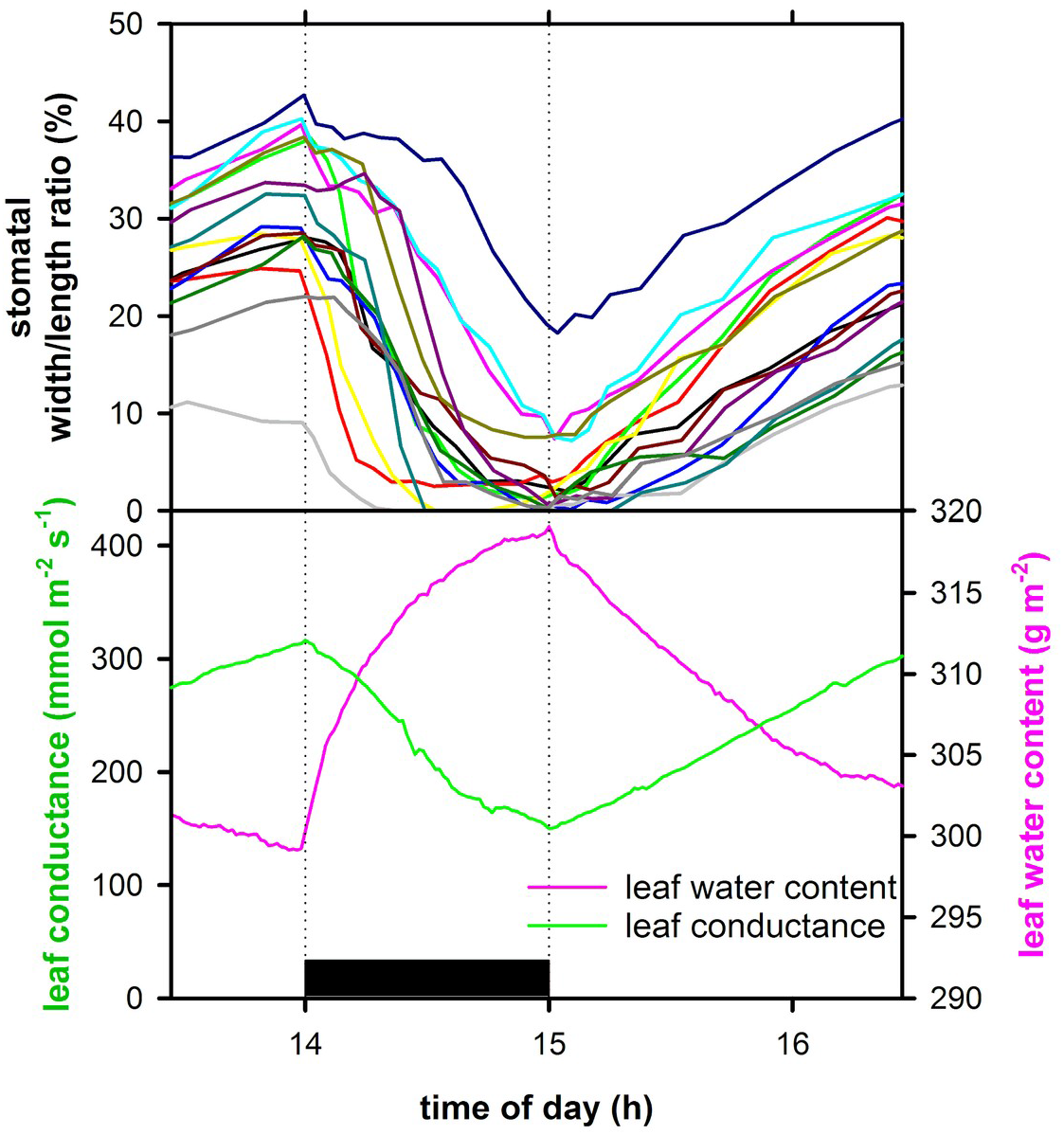
The typical dynamic response of leaf conductance, leaf water content and apertures of 15 microscopically observed stomata to intermittent darkening (1h). The experiment was performed at a temperature 24°C, a VPD of 1.05 kPa and a PPFD when illuminated of 450 µmol m^-2^ s^-1^.

### Experimental setup

The experiments presented here serve to demonstrate sensor performance and possible applications in combination with other methods, They were conducted with the LWC-sensor installed in a cuvette of a customized gas-exchange system (Walz GmbH, Effeltrich, Germany) allowing LWC measurements under controlled temperature light and VPD simultaneously with H_2_O gas-exchange measurements and microscopic observations of stomata on the abaxial leaf surface of the same leaf see (Kaiser and Legner 2007). This not only enables the transpiration based calibration procedure described above but also allows to calculate a continuous leaf water budget from measured transpiration and rate of change in leaf water content. The rate of influx through the petiole is the difference between efflux by transpiration and the rate of change in LWC.

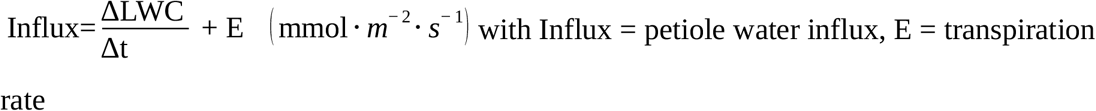

Hereby, a continuous determination of leaf water fluxes is possible.

The sensor was tested on leaves of *Vicia faba* L. grown in a greenhouse under ample supply of light water and nutrients. Variations in LWC were induced by excision of the leaf, darkening, changing VPD and gradual soil drying followed by sudden rewatering.

## Results and discussion

The sensor uses the IR-absorbtion ratio at 1450 nm vs 1050 nm to provide a precise real time measurement leaf water content. This ratiometric dual-beam transmission measurement using one water sensitive and one less water sensitive has been successfully applied previously (Sancho-Knapik et al. 2011; Cecilia et al. 2022) to record variations in leaf water status. The goal of the approach presented here was to overcome the problems of using only a limited spectral information on structurally complex and diverse objects with different biochemical compositions to estimate the content of water which is only one of several effector of changes in IR transmission. While the transmission ratio at the spectral bands around 1450 and 1050 nm has a strong relation to LWC content, the interfering influences of leaf structure and differences in biochemical composition preclude the possibility of a universal calibration valid across species and leaf morphologies. This is evident from the calibrations for maize and faba bean (fig. 2). The different calibration parameters for different growing conditions demonstrate that a calibration per species may not be sufficient, but that plants from different growing conditions require a separate calibration. The reasons for differences between calibrations are speculative. In the case of the Vicia faba, a 25% higher dry mass was found in the outdoor grown leaves, which might explain the differences, but they may also result from variations in internal leaf structure or reflective properties of the leaf surface. Irrespective of the reasons for variations in calibration parameters, the examples (fig. 2) demonstrate that a calibration for a batch of experimental plants from identical growing conditions can result in an accuracy of LWC estimation of c. 3-6%. In many experiments this accuracy can be sufficient especially if the focus is on detecting relative changes of LWC where precision is more important than accuracy. In the experiments presented here however, the simultaneous measurement of transpiration offered the opportunity to calibrate the sensor specifically for the leaf under investigation using cumulated transpirational water loss of the detached leaf together with gravimetric determination of the remaining LWC to achieve a both accurate and precise calibration. Calibrated in this way, the limited spectral information from a two-wavelength-measurement is sufficient to record water contents both precisely and accurately. It should be kept in mind, however, that the spot of LWC measurement is small and may not be fully representative for the whole leaf whose water content is determined by the integrating weight and transpiration measurements during calibration. Future technical improvement should address this problem by using more or wider measurement beams.

### Examples of applications

In order to test the usability of the sensor to detect fluctuations in LWC induced by changes in environmental conditions in different experimental settings, the sensor was installed into a gas-exchange cuvette with full control of temperature, VPD and Light and the option to measure transpiration and stomatal apertures simultaneously (Kaiser 2009). Fluctuations of LWC were induced by varying PPFD, VPD, cutting the petiole and by varying the soil water content.

Darkening a transpiring leaf will lead to a decrease of transpiration due to a quick drop in leaf temperature and subsequent stomatal closure. The changed flow balance between uptake through the petiole and transpiration will lead to an increase in LWC. The dynamics of this response was observed in detail with the LWC sensor (fig. 4) in combination with cuvette measurements of leaf conductance. Upon darkening LWC content increased by about 7%, approaching a new equilibrium after 1h. After re-illumination this process was reversed. The LWC sensor tracked these changes in high temporal and signal resolution which showed important details. For example, the change of LWC followed a typical relaxation kinetics with the rate of change being largest just after darkening/re-illumination and decreasing towards the new equilibrium. Together with the simultaneously measured stomatal responses of stomata and gas fluxes a dynamic in-situ analysis of leaf water relations is possible.

In another experiment the response to cutting a leaf was measured (fig. 5), demonstrating the classical Iwanoff-effect (Iwanoff 1928) of hydropassively opening of stomata upon the onset of water loss. This temporal opening is known to rely on the so called mechanical advantage of guard cells (De Michele and Sharpe 1973) which causes opening of stomatal pores after a turgor decline due to a negative water balance. In this experiment the LWC-sensor informed about the dynamics of leaf water content, showing that the maximum rate of water loss coincided with the maximum of leaf conductance 15 minutes after cutting the petiole. This demonstrates the action of a positive feedback-loop between ‘wrong way’ stomatal responses and leaf hydration (Cowan 1972) which initially led to an acceleration of water loss until active stomatal closure after 20 min was able to counteract and to close stomata.

**Fig. 5.**
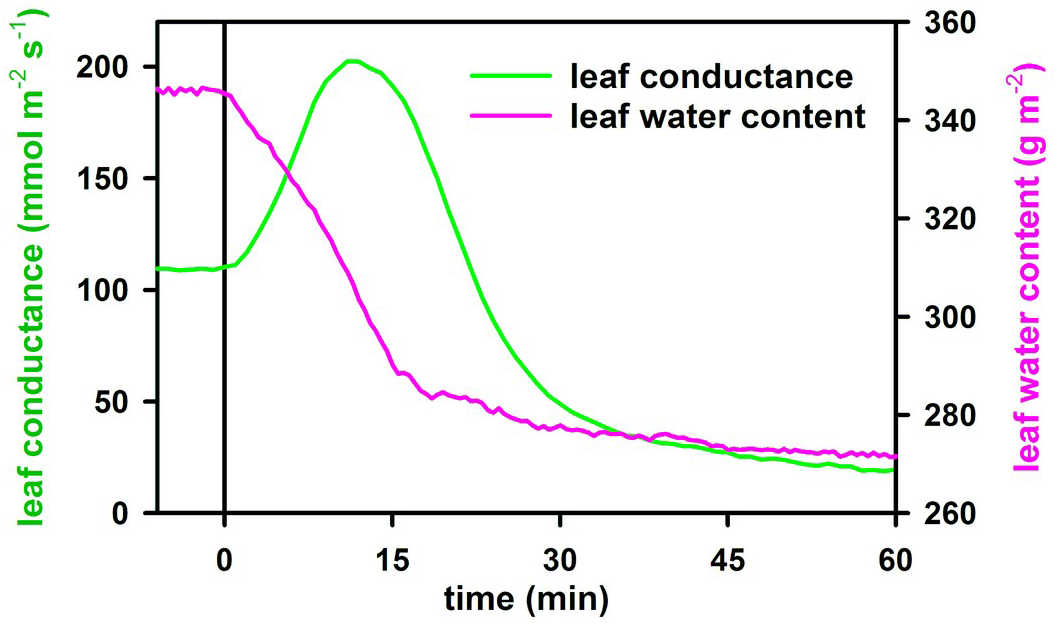
Response of LWC and leaf conductance to detachment of the leaf from the stem. The experiment was performed at a temperature of 24°C, a VPD of 1.05 kPa and a PPFD of 400 µmol m^-2^ s^-1^. Representative response of numerous recordings on detached leaves.

The measurement of a LWC response to a more temporal disturbance of leaf water status by a temporal increase of VPD from 1.1 kPa to 1.85 kPa (fig. 6) revealed a complex dynamic of LWC and its interaction with stomatal movements and gas exchange. After switching to high VPD, similar to the previous leaf cutting experiment, a temporal decline in LWC caused temporary hydropassive stomatal opening followed by active closure and consequently recovery of LWC to a value almost as high as in lower VPD. Switching back to low VPD revealed the inverse dynamic, an increase in LWC caused transient further stomatal closure. At this time, the positive feedback between leaf hydration, passive stomatal responses and transpiration supposedly lead to leaf water contents much higher than at the beginning. A delayed active reopening by increased transpiration restores the previous balance between water uptake into the leaf and transpiration and a recovery of the initial LWC. Notably, while short term LWC disturbances apparently were accentuated by hydraulic positive feedback, longer term stomatal adjustments recovered a similar LWC irrespective of VPD as would be expected from a feedback control of LWC. This suggests that the inclusion of LWC measurements into such experiments can provide the missing causal link between transpiration induced changes in leaf tissue hydration and stomatal feedback responses and could allow a more complete analysis of the interactions of hydraulic and pyhsiologic processes and their dynamic interaction.

**Fig. 6.**
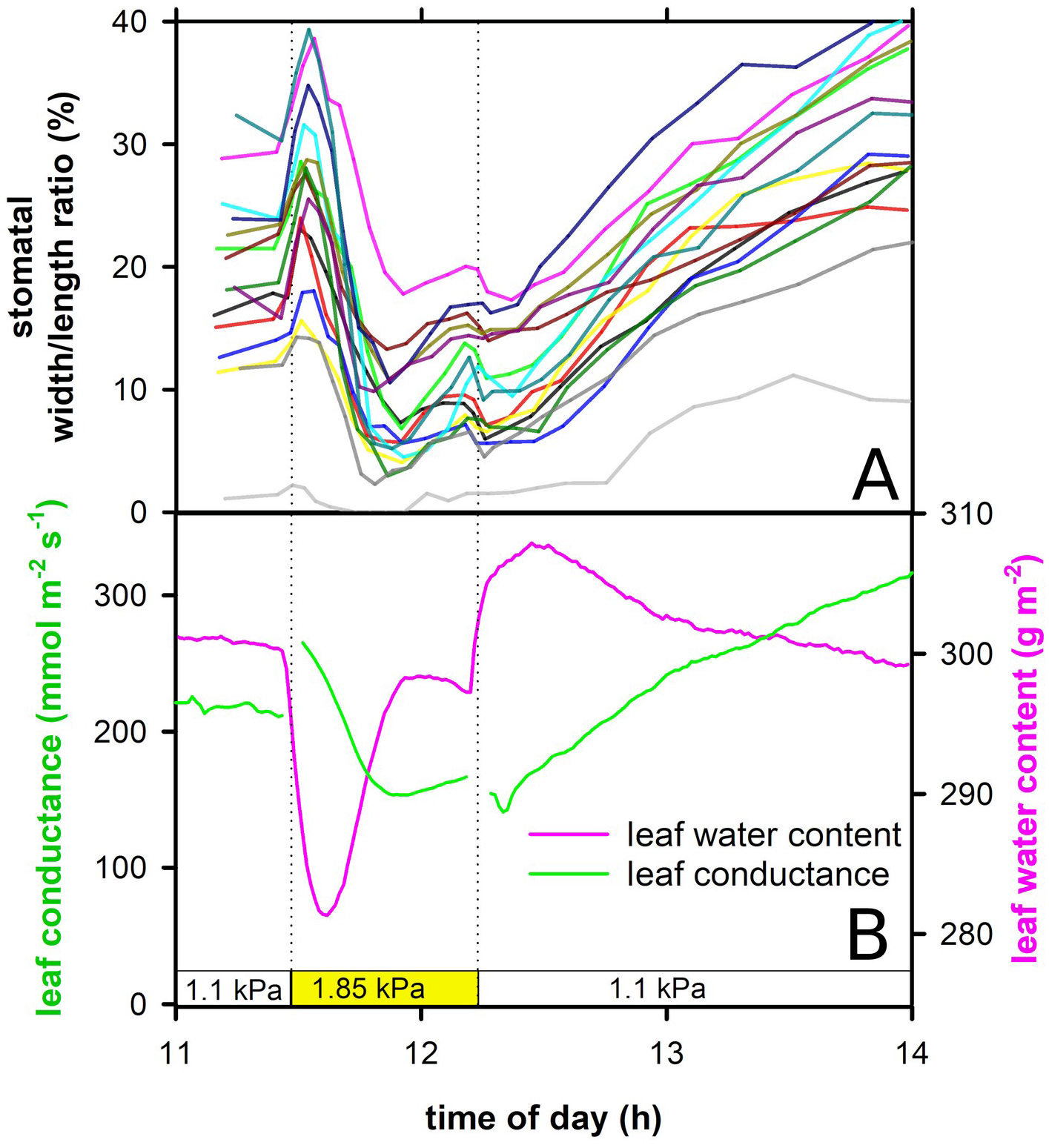
Typical responses of stomata, leaf conductance and LWC to a temporary increase (1h) of VPD from 1,1 kPa to 1,85 kPa. A) Microscopically observed apertures of 15 individual stomata. B) LWC and leaf conductance. The measurement of g_L_ was interrupted during transitions to a different VPD. The experiment was performed at a temperature of 25°C, and a PPFD of 400 µmol m^-2^ s^-1^.

The method combination of gas-exchange with LWC-measurement was also used in a long term soil drying and rewatering experiment (fig. 7 and 8), to assess whether the sensor provides stable and meaningful signals over terms of several days. During gradual decline in soil water content, LWC stayed largely constant for the first four days and only then started to decline. Stomatal conductance however started to decline earlier and was clearly reduced on the last two days of the experiment apart from a short temporary increase in the morning. Apparently stomatal responses at first successfully acted to preserve water. As LWC at this stage was not yet affected, this closure can be seen as ‘pre-emptive’. Only on day five LWC started to decrease at an increasing rate leading to a quick and visually apparent wilting process. At this point, by rewatering the plant, a recovery was initiated and monitored.

**Fig. 7.**
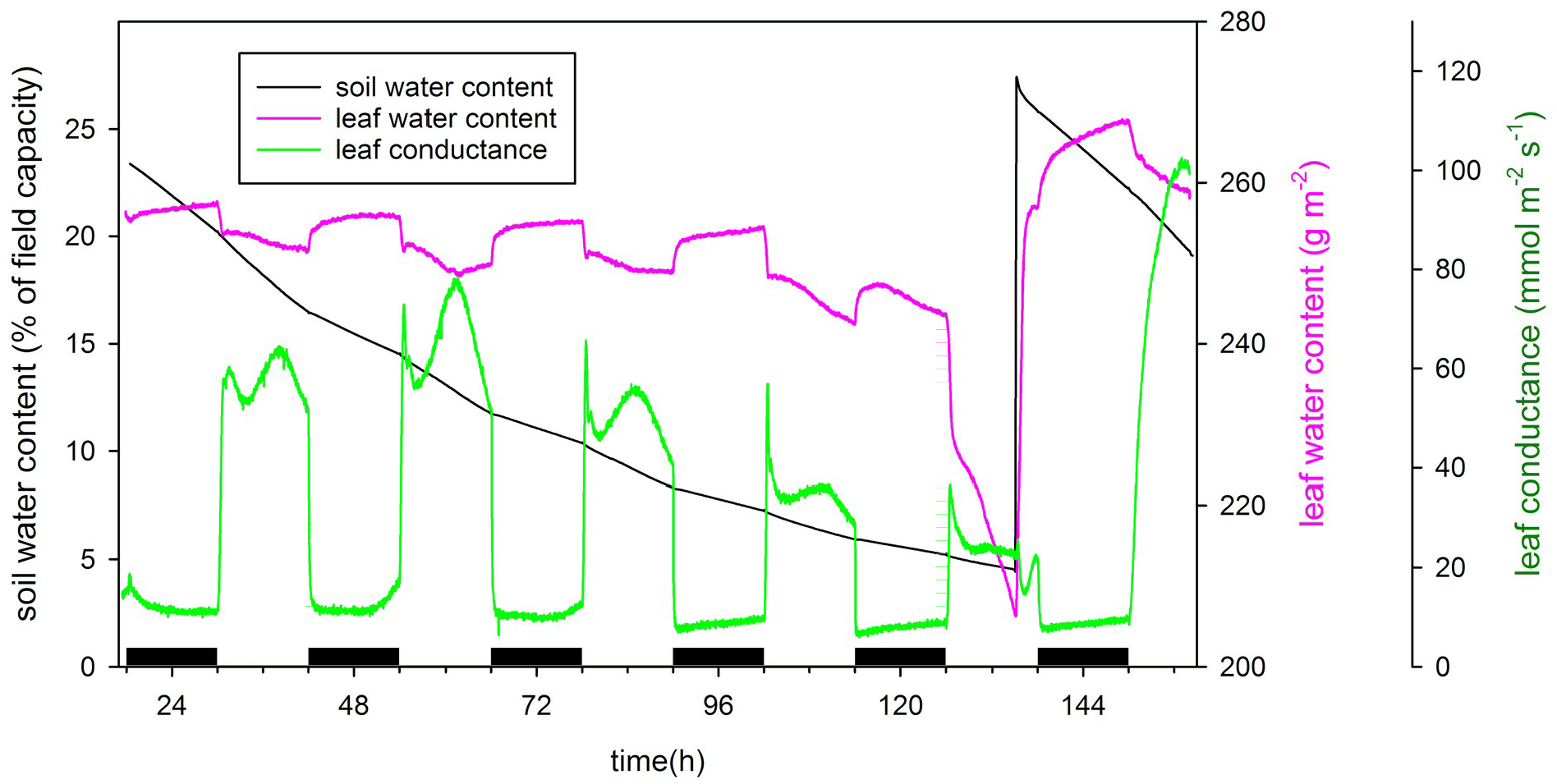
Leaf water content and leaf conductance during a soil drying and rewatering pot experiment on a leaf enclosed in a gas exchange chamber. Soil water content was measured by placing the pot on top of a continuously recording balance. The leaf was kept at 20°C, a VPD of 1.5 kPa and a 12/12 L/D cycle with a PPFD of 50 µmol m^-2^ s^-1^.

**Fig. 8.**
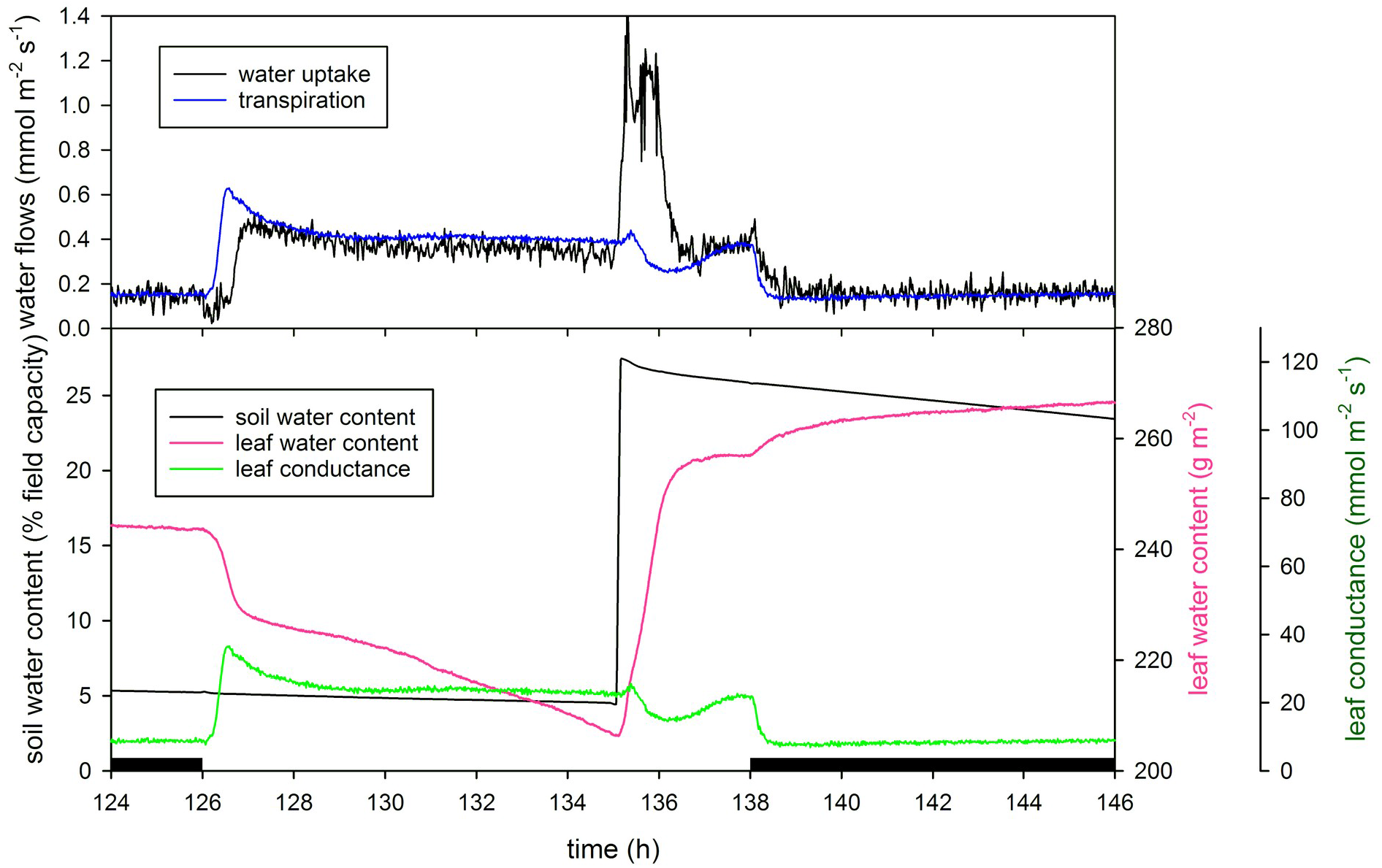
Leaf water content, leaf conductance and balance of water flows during wilting and re-watering of a leaf. Detail from the final stage of the soil drying experiment shown in fig. 7. The upper panel shows measured transpiration and water uptake through the petiole as calculated from the leaf water balance equation.

The dynamics during the final stage of wilting and the subsequent recovery here serve as an example, how a combination of LWC and transpiration measurements can be used to calculate water uptake through the petiole, thereby making up a continuous leaf water balance (fig 8). The comparison of leaf water uptake and transpiration showed that starting from the morning of the last day of the drought experiment, transpiration exceeded uptake, resulting in steady decline of LWC until wilting was reached. Rewatering after observation of visible wilting resulted in a rapid recovery completed within less than an hour. Noteworthy again is, that continuous observation of LWC and transpiration allows a flow rate determination even in these rapid transitory events which are governed by rapidly changing flow rates and leaf capacitance (Blackman and Brodribb 2011; Schymanski et al. 2013). Strikingly, LWC after recovery was higher than at the beginning of the experiment, which could be explained by osmotic adjustment during the 5 days of gradual drying resulting in increased osmotic potential and hence higher final water content. In *Vica faba*, changes in ion fluxes (Shabala et al. 2000) and in expression patterns of sugar metabolism related genes conforming to processes of osmotic adjustment (Ghouili et al. 2021) were found after exposure to osmotic stress. The reports on the actual existence and magnitude of osmotic adjustment, however, remain contradictory (Amede et al. 1999; Khallafallah et al. 2008; Abid et al. 2017). The LWC sensor in combination with other direct measurements of osmotic or total water potential appears as a promising tool to investigate the research field of osmotic adjustment.

To summarize the lessons learned from these example experiments, the presented method appears best suited for experiments requiring precise continuous recording of relative changes in LWC with sufficient SNR to detect even slight variations in near real time. The accuracy depends on the chosen calibration method and is high for a single leaf calibration, whereas the less accurate species specific calibration is sufficient for many applications and will maintain the high precision when measuring relative changes in LWC. The use of only two wavelengths thus means, that in order to achieve both high precision and accuracy the additional effort of a per leaf calibration procedures is required. The reward is however a small and simple sensor setup which does not impose many limitations on the choice of other simultaneous measurements. Here, combinations with cuvette based gas exchange measurements under control of gas composition, and light level together with in-situ microscopy are demonstrated. Combinations with other non-invasive optical methods like chlorophyll fluorescence probing and spectroscopic measurements are feasible.

In combination with recording of leaf gas exchange a real time balance of leaf water uptake and loss via transpiration can be calculated enabling e.g. an analysis of leaf water capacitances and flow resistances (Blackman and Brodribb 2011).

It might be questioned that LWC is a relevant parameter for leaf water relations at all. The significance of measures for water status has been under discussion (Jones 2007). Water potential, osmotic potential and turgor potential are linked and co-varying according to the water potential equation with changing leaf water content. Frequently, water potential is considered as the most important measure, as it ultimately determines the direction of water transport. For many physiological processes however, leaf water content appears to be a better determinant (Sinclair and Ludlow 1985). For example, LWC decline has been linked to inhibited Photosynthesis, (Kaiser 1987; Lawlor and Tezara 2009) and altered Chloroplast movement (Nauš et al. 2016). It should be considered here, that small variations in LWC typical for sub-stress fluctuations in water status will only have a relatively small impact on water potential, while directly affecting cell volume and turgor pressure. Changes in cell volume and turgor pressure are controversely discussed as properties being sensed by cellular mechanims and feeding back into processes regulation water status (e.g. activation of proton pumps, adjustment of osmotic potential, guard cell responses, activation of key enzymes of ABA synthesis (McAdam and Brodribb 2016; Sack et al. 2018; Zhang et al. 2018). If changes in cell volume are the relevant property, direct measurements of LWC are a useful experimental tool. Continuous non-invasive turgor measurements would also be desirable but they turned out to be difficult and attempts to use mechanical force sensors (Zimmermann *et al*., 2008) while providing continuous recordings of relative changes suffer from lack of absolute calibration and disrupt the local tissue water balance by blocking transpiration at the measured site. Instead of direct turgor measurements, water content can serve as a good proxy for turgor as long as cell volume and cell wall elasticity remain constant. Although drought can induce long term changes in cell wall elasticity (Martínez *et al*., 2007) over short term cell wall properties can be assumed to be constant resulting in a fairly constant turgor-volume relation which makes LWC a good proxy for turgor changes. Measurements of LWC are definitely meaningful when performed simultaneously with transpiration measurements, allowing a quantitative coverage of all liquid and gaseous water exchanges of a leaf.

## Abbreviations

LWC: leaf water content
NIR: near infrared
rH: relative humidity
SNR: signal to noise ratio
VPD: vapour pressure deficit

## Acknowledgements

I gratefully acknowledge the skillfull technical assistance of late Frank-Peter Rapp. Thanks go to Jon Henningsen for aid in calibrations.

## Data availability statement

Data are available on request from the author.

